# Identification of a Novel Osteogenetic Oligodeoxynucleotide (osteoDN) that Promotes Osteoblast Differentiation in a TLR9-Independent Manner

**DOI:** 10.1101/2022.03.21.485101

**Authors:** Yuma Nihashi, Mana Miyoshi, Koji Umezawa, Takeshi Shimosato, Tomohide Takaya

## Abstract

Recent studies have revealed that oligodeoxynucleotides (ODNs) designed from genome sequences have the potential to regulate cell fate. Currently, ODNs that conduct cell differentiation are nanomolecular drug candidates for regenerative medicine. Herein, we demonstrate that iSN40, an 18-base ODN derived from the *Lacticaseibacillus rhamnosus* GG genome, promoted the differentiation and calcification of osteoblasts that play a central role during bone formation. In the murine osteoblast cell line MC3T3-E1, iSN40 enhanced alkaline phosphatase activity at the early stage of differentiation and facilitated calcium deposition at the late stage by inducing the expression of osteogenic genes such as Msx2, osterix, collagen type 1α, osteopontin, and osteocalcin. Intriguingly, the CpG motif within iSN40 was not required for its osteogenetic activity, indicating that iSN40 functions in a TLR9-independent manner. These data suggest that iSN40, serving as an osteogenetic ODN (osteoDN), as a drug seed that target osteoblasts for bone regeneration.

## 1. Introduction

The bone is a metabolically active organ that remodels continuously throughout life. Skeletal homeostasis is maintained via bone remodeling through maintenance of balance between bone formation by osteoblasts and bone resorption by osteoclasts. Differentiation of osteoblasts into osteocytes is an essential step during bone formation, which is regulated by the osteoblast-specific transcription factors, Runx2 and osterix, promoting the synthesis, deposition, and mineralization of the extracellular matrix. However, aging impairs osteoblast activity such as the expression of collagen type 1α [1], which is considered the main pathological reason for the decline in bone mass, typically osteoporosis, in elderly people [2]. Therefore, activation of osteoblasts is one of the key strategies to realize bone regeneration. Several agents have been developed to target osteoblast-oriented bone formation in osteoporosis therapy [3]. Parathyroid hormone and its N-terminal fragment, teriparatide, activate bone anabolism and increase bone density by directly targeting osteoblasts [4]. Osteoblasts secrete receptor activator of NF-κB ligand (RANKL), that is recognized by osteoclasts, and osteoclastogenesis is initiated. Denosumab, a specific decoy receptor for RANKL, increases bone mineral density and prevents bone resorption [5]. Currently, denosumab is the most preferred treatment for post-menopausal osteoporosis [6]. Sclerostin inhibits canonical Wnt/β-catenin signaling and increases RANKL production in osteocytes [7]. Thus, romosozumab, a sclerostin antibody, activates the Wnt pathway to stimulate bone formation and suppress resorption [8]. The beneficial effects of these hormones and antibodies in osteoporosis therapy have been robustly proven. However, with the current technology, these protein/peptide-based drugs are expensive and unstable to be produced in large quantities for the treatment of a large number of patients with osteoporosis worldwide.

As nucleic acids offer several advantages, such as chemical synthesis, low-cost manufacturing, and stability during storage, they are potential nanomolecules for next-generation drugs. A wide variety of nucleotides have been clinically applied; antisense nucleotides that regulate gene expression [9], aptamers that target proteins [10], and ligands of Toll-like receptors (TLRs) that modulate the innate immune system [11] can specifically access their diverse targets. Furthermore, although there are only few studies available, oligodeoxynucleotides (ODNs) from genome sequences have been reported to regulate stem cell fate, and may be employed for regenerative medicine. A 27-base cytosine (C)- rich ODN designed from the human mitochondrial genome, named MT01 ([ACC CCC TCT]_3_), was initially identified as an immunosuppressive ODN that inhibits the proliferation of human peripheral blood mononuclear cells [12]. Intriguingly, MT01 promoted the proliferation and differentiation of the human osteoblast-like cell line MG63 [13], rat bone marrow mesenchymal stem cells [14], and murine osteoblast cell line MC3T3-E1 [15] by upregulating RUNX2 phosphorylation via activation of the ERK/MAPK pathway [16]. Thus, MT01 treatment reduced alveolar bone loss in rat periodontitis in vivo [14]. These studies suggest that genomic ODNs could be used as drug seeds for bone regeneration.

We recently screened an 18-base ODN library designed from the genome sequence of the lactic acid bacterium *Lacticaseibacillus rhamnosus* GG [17] to determine its effect on myogenic differentiation. The 18-base guanine (G)-rich ODN, iSN04, promoted differentiation and suppressed inflammation in myogenic precursor cells called myoblasts [18-22]. iSN04, termed myogenetic ODN (myoDN), formed a globular structure of 1-nm radius and served as an anti-nucleolin aptamer in a TLR-independent manner [18]. The discovery of myoDN demonstrates that bacterial genome sequences are promising platform for providing novel ODNs that can control cell differentiation. Herein, we report an osteogenetic ODN (osteoDN) that facilitates the differentiation and calcification of osteoblasts, which was identified from the same ODN library used for myoDN study. This study provides new insights into drug development for bone regeneration.

## 2. Materials and Methods

### 2.1. Chemicals

ODN sequences used in this study are listed in Table S1. Phosphorothioated (PS)-ODNs and 6-carboxyfluorecein (6-FAM)-conjugated PS-ODNs were synthesized and purified using HPLC (GeneDesign, Osaka, Japan). PS-ODNs and ascorbic acid (AA) (Fujifilm Wako Chemicals, Osaka, Japan) were dissolved in endotoxin-free water. An equal volume of solvent was used as negative control.

### 2.2. Cell Culture

MC3T3-E1 cell line (RCB1126) was provided by the RIKEN BRC (Tsukuba, Japan) through the Project for Realization of Regenerative Medicine and the National Bio-Resource Project of the MEXT, Japan. The cells were cultured at 37°C with 5% CO_2_ throughout the experiments. Undifferentiated MC3T3-E1 cells were maintained in a growth medium (GM) consisting of α-MEM (Nacalai, Osaka, Japan), 10% fetal bovine serum (FBS) (HyClone; GE Healthcare, UT, USA), and a mixed solution of 100 units/ml penicillin and 100 μg/ml streptomycin (P/S) (Nacalai). To induce osteogenic differentiation, confluent MC3T3-E1 cells were cultured in a differentiation medium (DM) consisting of α-MEM, 10% FBS, 10 nM dexamethasone (Fujifilm Wako Chemicals), 10 mM β-glycerophosphate (Nacalai), and P/S. To facilitate differentiation, 15 or 50 μg/ml AA was added to GM or DM as required.

### 2.3. Alkaline Phosphatase (ALP) Staining

For screening, MC3T3-E1 cells were seeded on 96-well plates (1.0×10^4^ cells/100 μl GM/well) for screening or on 12-well plates (1.0×10^4^ cells/well) for high-resolution imaging. The next day, the medium was replaced with GM or DM containing 10 μM PS-ODNs. After 48 h, ALP enzymatic activity of the cells was visualized using ALP Stain Kit (Fujifilm Wako Chemicals) according to the manufacturer’s instruction. Phase-contrast images were obtained using EVOS FL Auto microscope (AMAFD1000; Thermo Fisher Scientific, MA, USA). ALP-positive area was quantified using ImageJ software (National Institute of Health, USA).

### 2.4. Alizarin Staining

MC3T3-E1 cells were seeded on 24-well plates (3.0-5.0×10^4^ cells/well) and cultured in GM until they become confluent. The medium was then replaced with DM containing ODNs every 3-4 days. The cells were fixed with 2% paraformaldehyde and stained with 1% w/v alizarin red S solution (Fujifilm Wako Chemicals). Bright-field images were captured using EVOS FL Auto microscope. Alizarin-positive area was quantified using ImageJ software.

### 2.5. Quantitative Real-Time RT-PCR (qPCR)

MC3T3-E1 cells were seeded on 60-mm dishes (3.0×10^5^ cells/dish). The next day, the medium was replaced with GM or DM containing 10 μM iSN40. Total RNA was isolated using NucleoSpin RNA Plus (Macherey-Nagel, Düren, Germany) and reverse transcribed using ReverTra Ace qPCR RT Master Mix (TOYOBO, Osaka, Japan). qPCR was performed using GoTaq qPCR Master Mix (Promega, WI, USA) with the StepOne Real-Time PCR System (Thermo Fisher Scientific). The amount of each transcript was normalized to that of the 3-monooxygenase/tryptophan 5-monooxygenase activation protein zeta gene (*Ywhaz*) and presented as fold-changes. The primer sequences are listed in Table S2.

### 2.6. Trivial Trajectory Parallelization of Multicanonical Molecular Dynamics (TTP-McMD)

Starting with the simulation of iSN40 and MT01 structures built from their DNA sequences using NAB in AmberTools [23], an enhanced ensemble method, TTP-McMD [24], was used to sample the equilibrated conformations at 310 K. In the TTP-McMD, energy range of the multicanonical ensemble covered a temperature range from 280 K to 380 K. Sixty trajectories were used, and the production run was conducted for 40 ns in each trajectory (total 2.4 μs). Throughout the simulation, amber ff12SB force field [25] was used, whereas the solvation effect was represented by the generalized-born model [26].

### 2.7. Statistical Analysis

The results are presented as the mean ± standard error. Statistical comparisons between two groups were performed using unpaired two-tailed Student’s *t*-test and among multiple groups using one-way analysis of variance followed by Scheffe’s *F*-test. Statistical significance was set at *p* < 0.05.

## 3. Results

### 3.1. iSN40 Promotes Osteoblast Differentiation

Forty-four 18-base PS-ODNs designed from the *Lacticaseibacillus rhamnosus* GG genome (iSN04 and iSN08-iSN50) were administered to MC3T3-E1 cells. In addition, two immunomodulatory PS-ODNs were concomitantly tested: CpG-2006, a TLR9 ligand that initiates inflammatory cascades [27], and Tel-ODN, a human telomeric ODN that suppresses immunological reactions depending on TLR3/7/9 [28]. The osteogenic effects of these PS-ODNs were investigated by measuring ALP enzymatic activity in the cells, a standard marker of osteoblast differentiation (Figure 1A). iSN40 (GGA ACG ATC CTC AAG CTT) markedly increased ALP-positive area to the same extent as AA, positive control for promoting osteogenic differentiation (Figure 1B). However, other PS-ODNs did not enhance ALP activity, indicating sequence-dependent osteogenetic activity of iSN40. iSN40-induced ALP activity was reproducibly confirmed by high-resolution images of the experiment performed independently of the screening (Figure S1). These results demonstrate that iSN40 serves as an osteoDN that promotes osteoblast differentiation.

**Figure 1.**
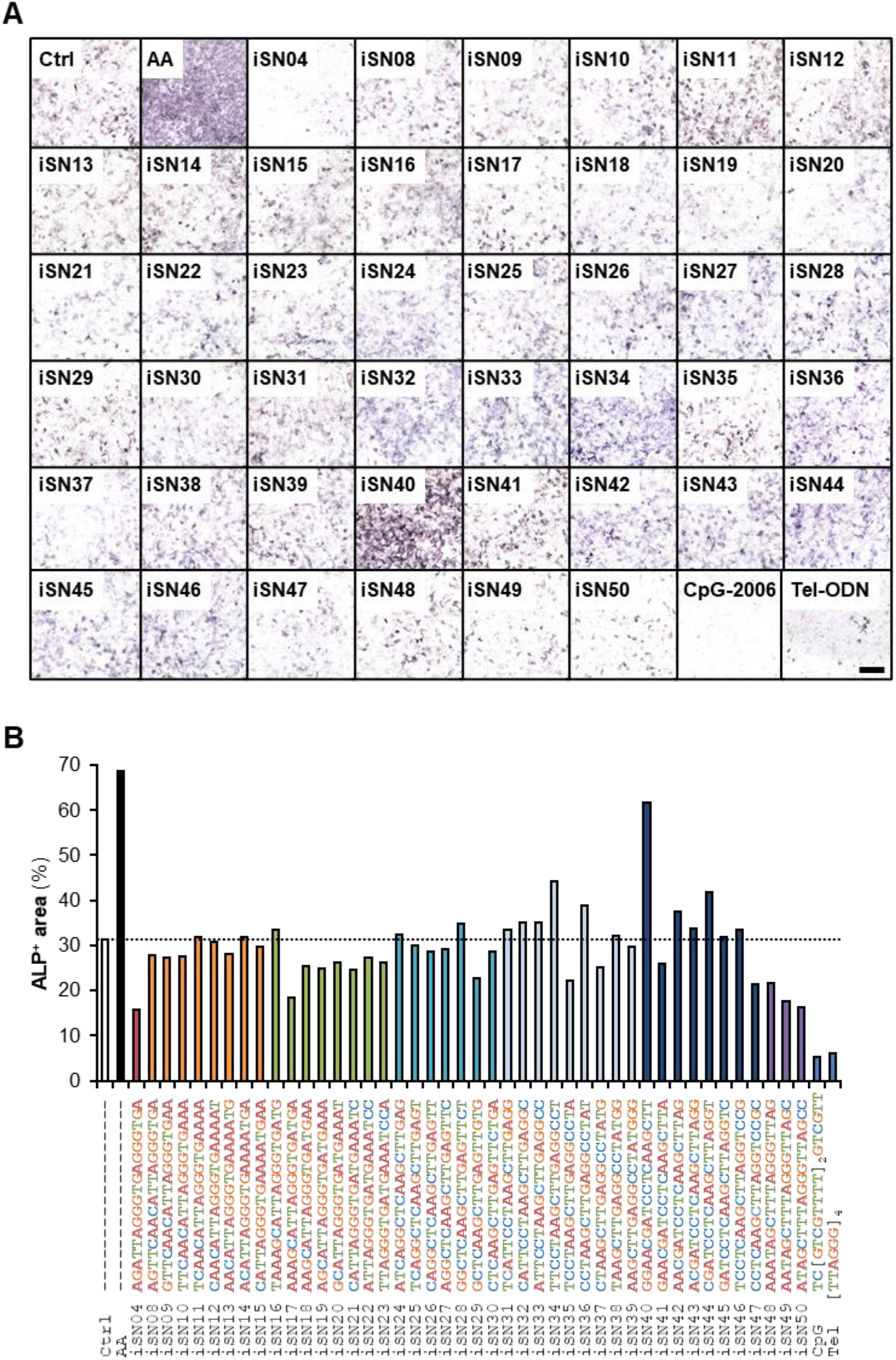
Identification of iSN40 as an osteoDN. (**A**) ALP staining images of MC3T3-E1 cells treated with 10 μM PS-ODNs in GM for 48 h. Scale bar, 200 μm. (**B**) Quantification of ALP-positive area.

### 3.2. iSN40 Modulates Osteogenic Gene Expression

The effect of iSN40 on osteogenic gene expression in MC3T3-E1 cells was investigated by qPCR. To analyze the early stage of differentiation, MC3T3-E1 cells were treated with iSN40 in GM containing AA for 24-48 h (Figure 2A). iSN40 increased the mRNA levels of Msx2, a homeobox transcription factor that promotes osteogenesis through bone morphogenic protein 2-induced signaling. In contrast, iSN40 did not alter the levels of Runx2, a master regulator that determines osteoblast lineage and inhibits bone maturation [29]. Interestingly, iSN40 significantly upregulated the expression of osterix (*Sp7*), a downstream target of Runx2. iSN40 did not improve mRNA levels of immature myoblast markers, collagen type 1α (*Col1a1*) and osteopontin (*Spp1*); however, iSN40 markedly induced osteocalcin (*Bglap2*), a hormone released from the bone [30]. To investigate the late stage of differentiation, MC3T3-E1 cells were treated with iSN40 in DM containing AA for 4-8 days (Figure 2B). iSN40 significantly elevated the levels of Msx2 on day 4 but not that of Runx2. Notably, by day 8, iSN40 upregulated the expression of osterix, collagen type 1α, osteopontin, and osteocalcin, which act downstream of Runx2 [31]. Compared with the results at the early stage, the effect of iSN40 on these gene expression was more significant at the late stage, suggesting a long-term effect of iSN40. These data indicate that iSN40 facilitates osteoblast differentiation by modulating osteogenic gene expression through the transcription factor Msx2 rather than Runx2.

**Figure 2.**
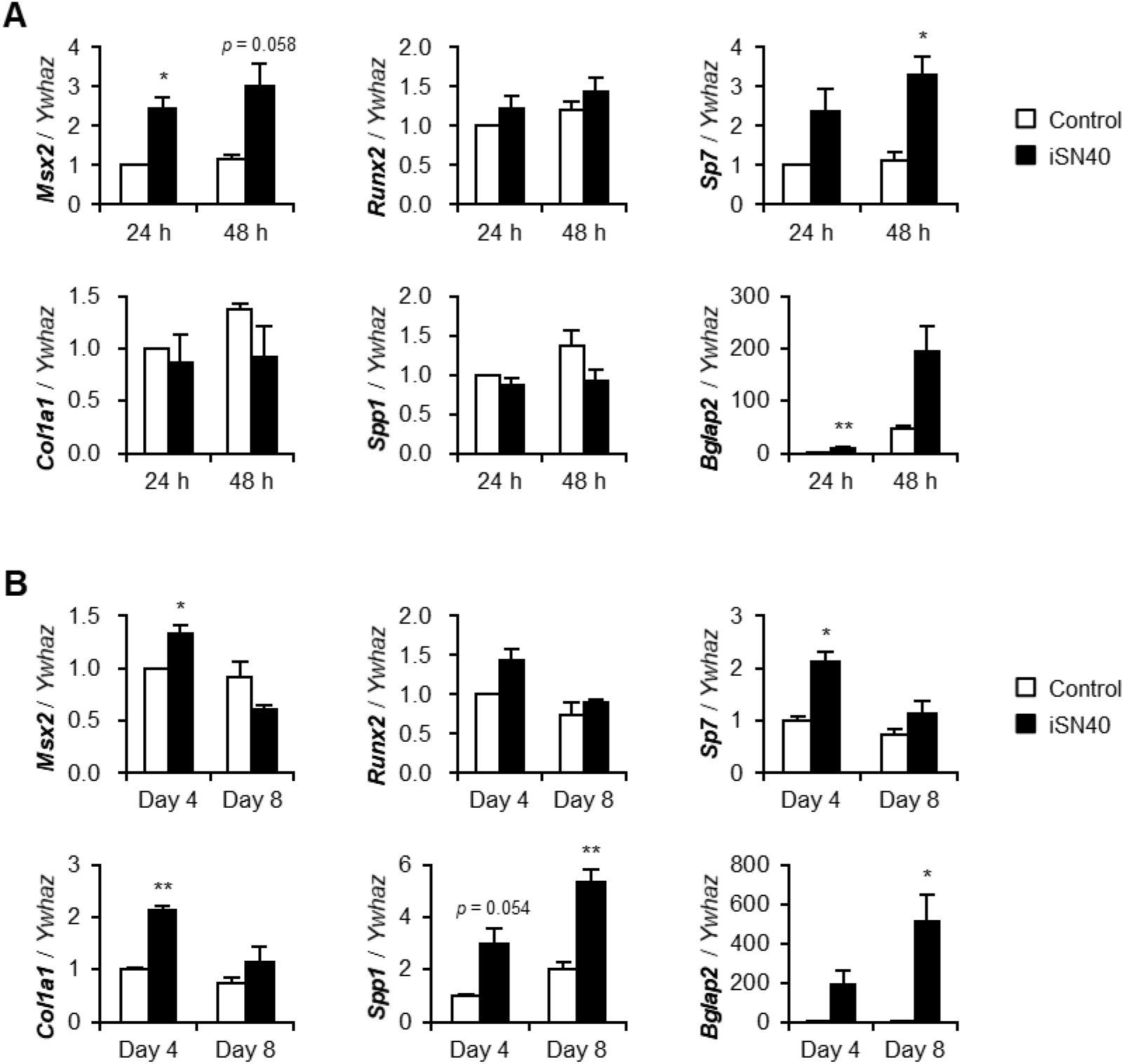
iSN40 induces osteogenic gene expression in MC3T3-E1 cells. (**A**) qPCR results of MC3T3E1 cells treated with 10 μM iSN40 in GM with 15 μg/ml AA for 24 or 48 h. (**B**) qPCR results of MC3T3-E1 cells treated with 10 μM iSN40 in DM with 15 μg/ml AA for 4 or 8 days. * *p* < 0.05, ** *p* < 0.01 vs control at each time point (Student’s *t* test). *n* = 3.

### 3.3. iSN40 Promotes Osteoblast Calcification

To investigate the effect of iSN40 on osteoblast calcification, MC3T3-E1 cells were treated with iSN40 in DM with or without AA for 12 days and then subjected to alizarin staining to visualize calcium deposition on the cells (Figure 3A). Alizarin-positive area was significantly increased by 10 μM iSN40 rather than 15 μg/ml AA. Moreover compared with individual treatments, co-treatment with iSN40 and AA further facilitated osteoblast calcification, indicating that iSN40 and AA synergistically promoted osteoblast mineralization. Although iSN40 was administered at a concentration of 10 μM in the above experiments, the dose response analysis revealed that 1 μM iSN40 was sufficient to induce osteoblast calcification in the presence of 15 μg/ml AA (Figure 3B). We used phosphorothioated iSN40 (PS-iSN40) in this studyto avoid degradation by nucleases, whereas native iSN40 did not promote calcification even in the presence of 50 μg/ml AA (Figure S2). This demonstrates that phosphorothioation is necessary for iSN40 to function as an osteoDN.

**Figure 3.**
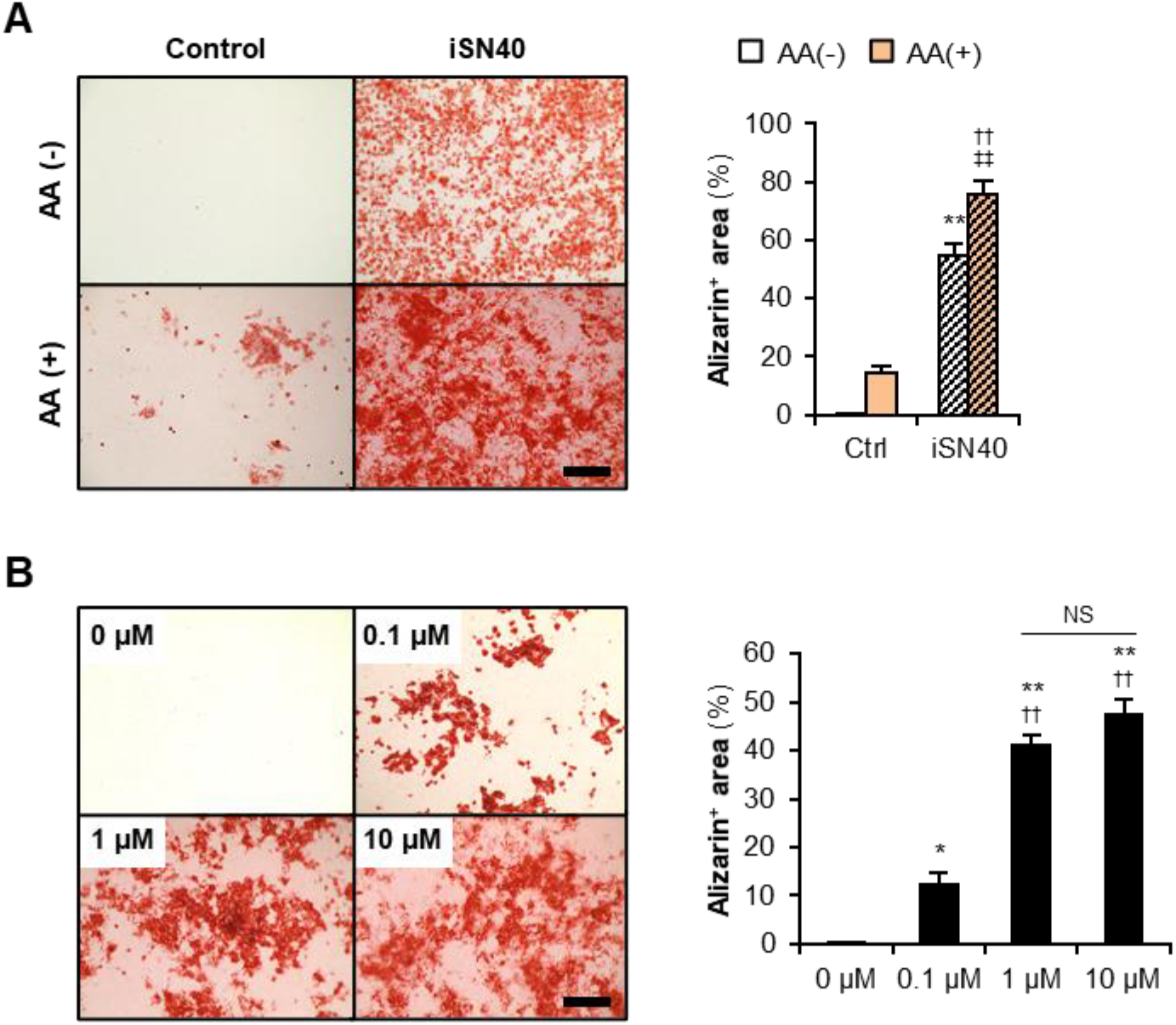
iSN40 induces calcification of MC3T3-E1 cells. (**A**) Representative images and quantification of alizarin staining of MC3T3-E1 cells treated with 10 μM iSN40 in DM with or without 15 μg/ml AA for 12 days. Scale bar, 200 μm. ** *p* < 0.01 vs control-AA(-), ^††^ *p* < 0.01 vs control-AA(+), ^‡‡^ *p* < 0.01 vs iSN40-AA(-) (Scheffe’s *F*-test). *n* = 4. (**B**) Representative images and quantification of alizarin staining of MC3T3-E1 cells treated with 0.1, 1, or 10 μM iSN40 in DM with 15 μg/ml AA for 9 days. Scale bar, 200 μm. * *p* < 0.05, ** *p* < 0.01 vs 0 μM; ^††^ *p* < 0.01 vs 0.1 μM; NS, not significant (Scheffe’s *F*-test). *n* = 4.

### 3.4. Osteogenetic Action of iSN40 is TLR9-Independent

iSN40 (GGA A<u>CG</u> ATC CTC AAG CTT) contained a CpG motif. It has been well studied that ODNs possessing unmethylated CpG motifs (CpG-ODNs) serve as TLR9 ligands and initiate innate immune system-induced inflammatory responses [32]. To examine the impact of the CpG motif within iSN40 on its osteogenetic activity, iSN40-GC (GGA A<u>GC</u> ATC CTC AAG CTT) was constructed, in which the CpG motif was substituted with GC. iSN40-GC enhanced ALP activity and calcification of MC3T3-E1 cells to the same extent as iSN40 (Figure 4). Conversely, a well-known TLR9 ligand, CpG-2006 [27], did not facilitate osteoblast mineralization (Figure 4B), which was further confirmed through ALP staining results (Figures 1 and S1). In addition, RT-PCR revealed that MC3T3-E1 cells do not express TLR9 (Figure S3), corresponding to the previous study [33]. These results demonstrate that CpG motif- and TLR9-independent osteogenetic action of iSN40.

**Figure 4.**
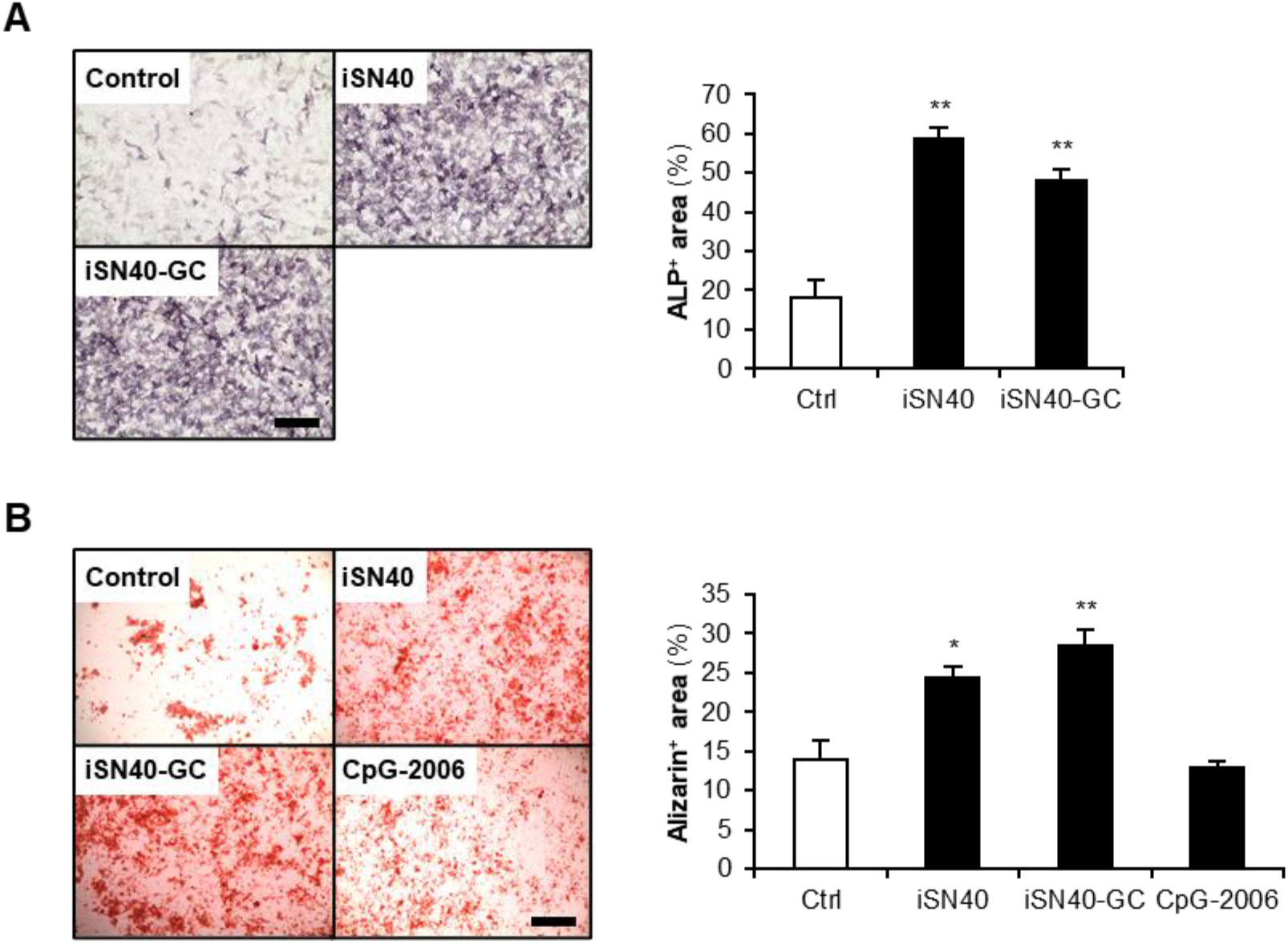
Action of iSN40 is TLR9-independent. (**A**) Representative images and quantification of ALP staining of MC3T3-E1 cells treated with 10 μM of iSN40 or iSN40-GC in DM with 50 μg/ml AA for 48 h. Scale bar, 100 μm. ** *p* < 0.01 vs control (Scheffe’s *F*-test). *n* = 4. (**B**) Representative images and quantification of alizarin staining of MC3T3-E1 cells treated with 1 μM PS-ODNs in DM with 50 μg/ml AA for 9 days. Scale bar, 200 μm. * *p* < 0.05, ** *p* < 0.01 vs control (Scheffe’s *F*-test). *n* = 4.

## 4. Discussion

The present study identified that iSN40, which is an 18-base ODN derived from the genome sequence of lactic acid bacteria, serves as an osteoDN that promotes differentiation and mineralization of osteoblasts, and it could be a potential nucleic acid drug candidate for bone regeneration via osteoblast activation. Although direct target and action mechanism of iSN40 are still unknown, the study results revealed that the osteogenetic function of iSN40 was TLR9-independent. The CpG motif within iSN40 was not required for its activity; A validated TLR9 ligand, CpG-2006 [27], enhanced neither ALP activity nor calcification of MC3T3-E1 cells; moreover, TLR9 was not expressed in MC3T3-E1 cells. TLR9-independent effect of iSN40 is crucial for its application in osteoporosis therapy. For example, CpG-ODNs generally upregulate interleukin (IL)-6 production via TLR9 [11]. IL-6 is a pro-osteoclastogenic cytokine [2] that is highly produced in aged osteoblasts [34]. The innate immune system, including IL-6 secretion, is closely related to bone homeostasis [35]; however, iSN40 is anticipated to activate osteoblasts without perturbing the microenvironment.

iSN40 upregulated the expression of osteogenic genes such as osterix, collagen type Iα, osteopontin, and osteocalcin, which are induced by Runx2 [31]. However, iSN40 did not alter Runx2 expression. The transcriptional activity of Runx2 is regulated via post-transcriptional modifications including phosphorylation, acetylation, sumoylation, and ubiquitination, which are mediated by numerous factors [36]. iSN40 may target such modulator proteins to enhance Runx2 activity. For instance, another osteoDN, MT01, has been reported to enhance Runx2 phosphorylation via activation of ERK and p38 MAPK [16], which potentiates the Runx2/osterix transcriptional machinery [37]. The effect of iSN40 on Runx2 protein needs to be examined in future studies.

iSN41-iSN47 are PS-ODNs analogous to iSN40; however, they did not enhance ALP activity of MC3T3-E1 cells. When iSN40 operates as an antisense nucleotide, iSN41-iSN47 would present partial osteogenetic activities. To discuss the mechanism of action of iSN40, a previous study reporting iSN04 is helpful. iSN04 is designed from the lactic acid bacteria, genome as same as iSN40, and serves as a myoDN that promotes myoblast differentiation in a TLR-independent manner [18]. iSN04 is taken up into cytoplasm without any carriers, forms a G-stacking structure, and physically interacts with nucleolin to interfere its function, indicating that iSN04 is an anti-nucleolin aptamer [18]. Thus, it is sufficiently possible that iSN40 works as an aptamer for target proteins. Administration of 6-FAM-conjugated iSN40 and iSN40-GC to MC3T3-E1 cells showed that they were autonomously incorporated into the cytoplasm within 30 min (Figure 5), suggesting that iSN40 acts intracellularly, not on the plasma membrane.

**Figure 5.**
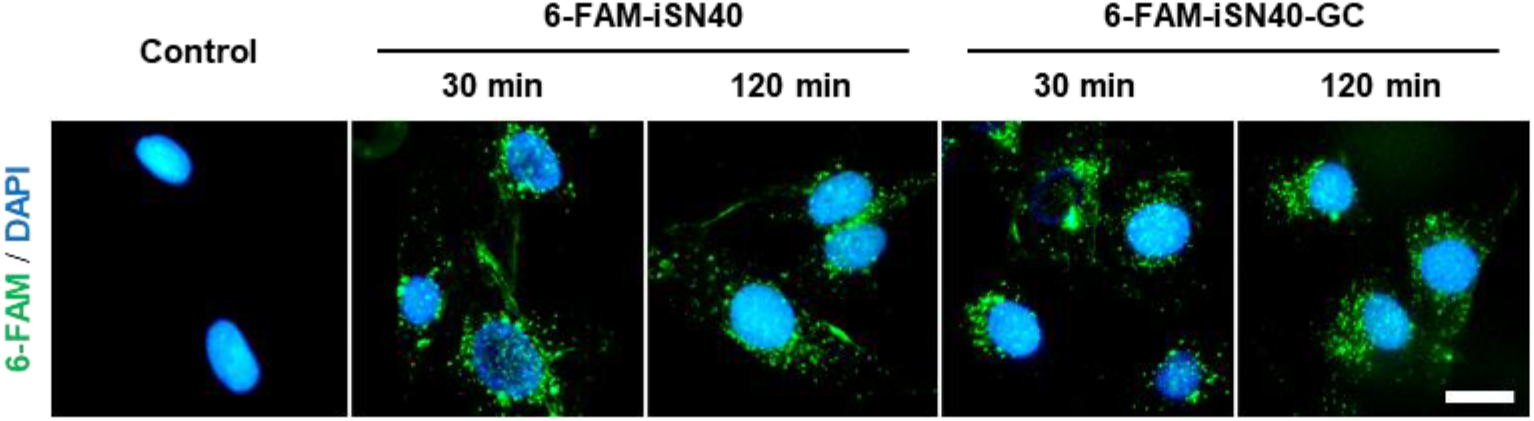
Intracellular incorporation of iSN40. Representative fluorescent images of MC3T3-E1 cells treated with 5 μg/ml 6-FAM-iSN40 and 6-FAM-iSN40-GC in GM for 30 or 120 min. Scale bar, 25 μm.

To further discuss the potential of iSN40 as an aptamer, conformation of iSN40 in water at 310 K was computationally simulated using the TTP-McMD (Figure 6A). iSN40 exhibited a compact globular structure (average radius: 1.01 nm) similar to that of iSN04 [18]. Interestingly, the predicted structure of another osteoDN, MT01, was substantially different from that of iSN40 (Figure 6B). As MT01 has a linear structure, the mechanisms of action of the two osteoDNs, iSN40 and MT01, are likely to be dissimilar. Indeed, iSN40 did not alter the mRNA levels of Runx2, but MT01 significantly induced Runx2 transcription [13,14]. To determine whether iSN40 is an aptamer, the iSN40-binding protein in osteoblasts needs to be identified. This would further promote the development and application of iSN40 as an osteoDN for bone regeneration.

**Figure 6.**
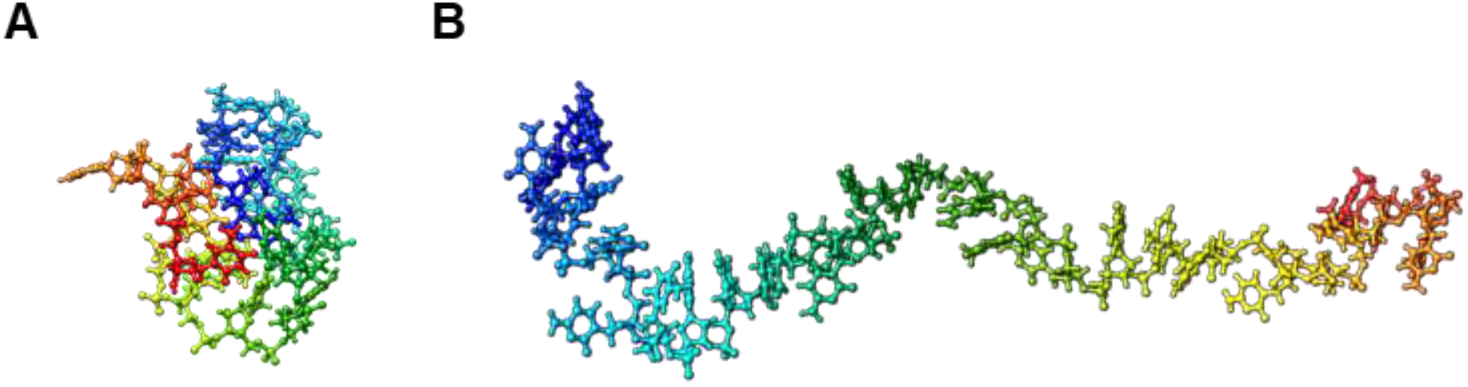
The most thermodynamically stable conformations of the ODNs in water at 310K simulated by the TTP-McMD. (**A**) iSN40. (**B**) MT01.

## 5. Conclusions

The present study identified that iSN40, an 18-base ODN designed from the lactic acid bacteria genome, serves as a novel osteoDN that promotes the differentiation and calcification of osteoblasts by modulating osteogenic gene expression in a TLR9-independent manner. iSN40 directly targets and activates osteoblasts, and will be useful in osteoporosis therapy as a nucleic acid drug that promotes bone formation and regeneration.

## 6. Patents

Shinshu University has been assigned the invention of osteoDN by T.T., Y.N., K.U., and T.S., and Japan Patent Application 2021-122713 has been filed on July 27, 2021.

## Author Contributions

Y.N. and T.T. designed the study. T.T. wrote the manuscript. Y.N. and M.M. performed experiments and data analyses. K.U. calculated the ODN structures. T.S. designed and provided the ODN library. All authors have read and agreed to the published version of the manuscript.

## Data Availability Statement

The raw data supporting the conclusions of this article will be made available by the authors, without undue reservation.

## Supplementary Materials

**Supplementary Figure S1.**
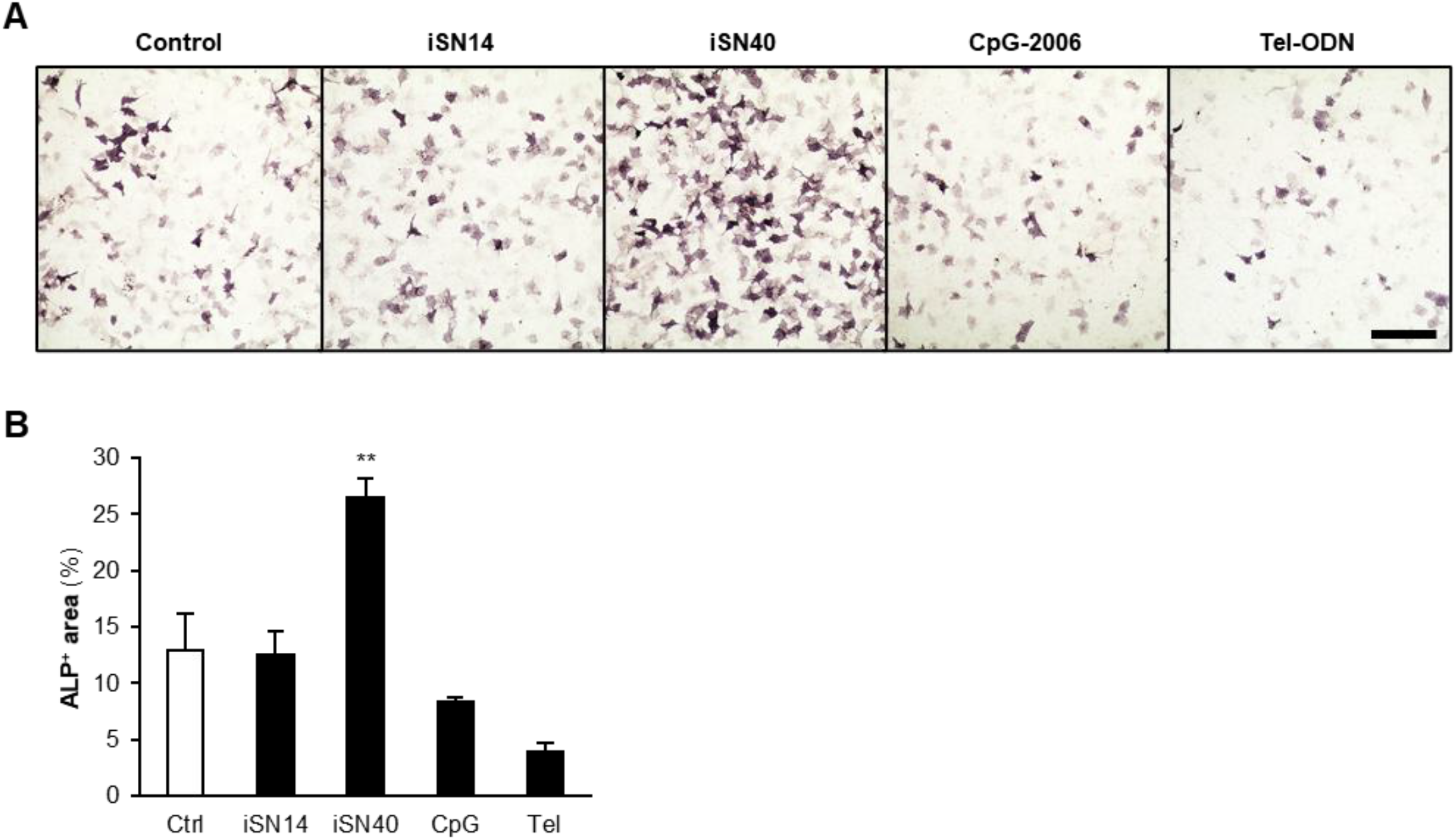
iSN40 enhances ALP activity in MC3T3-E1 cells. (**A**) Representative images of ALP staining of MC3T3-E1 cells treated with 10 μM PS-ODNs in GM for 48 h. Scale bar, 100 μm. (**B**) Quantification of ALP-positive area. ** *p* < 0.01 vs control (Scheffe’s *F*-test). *n* = 5.

**Supplementary Figure S2.**
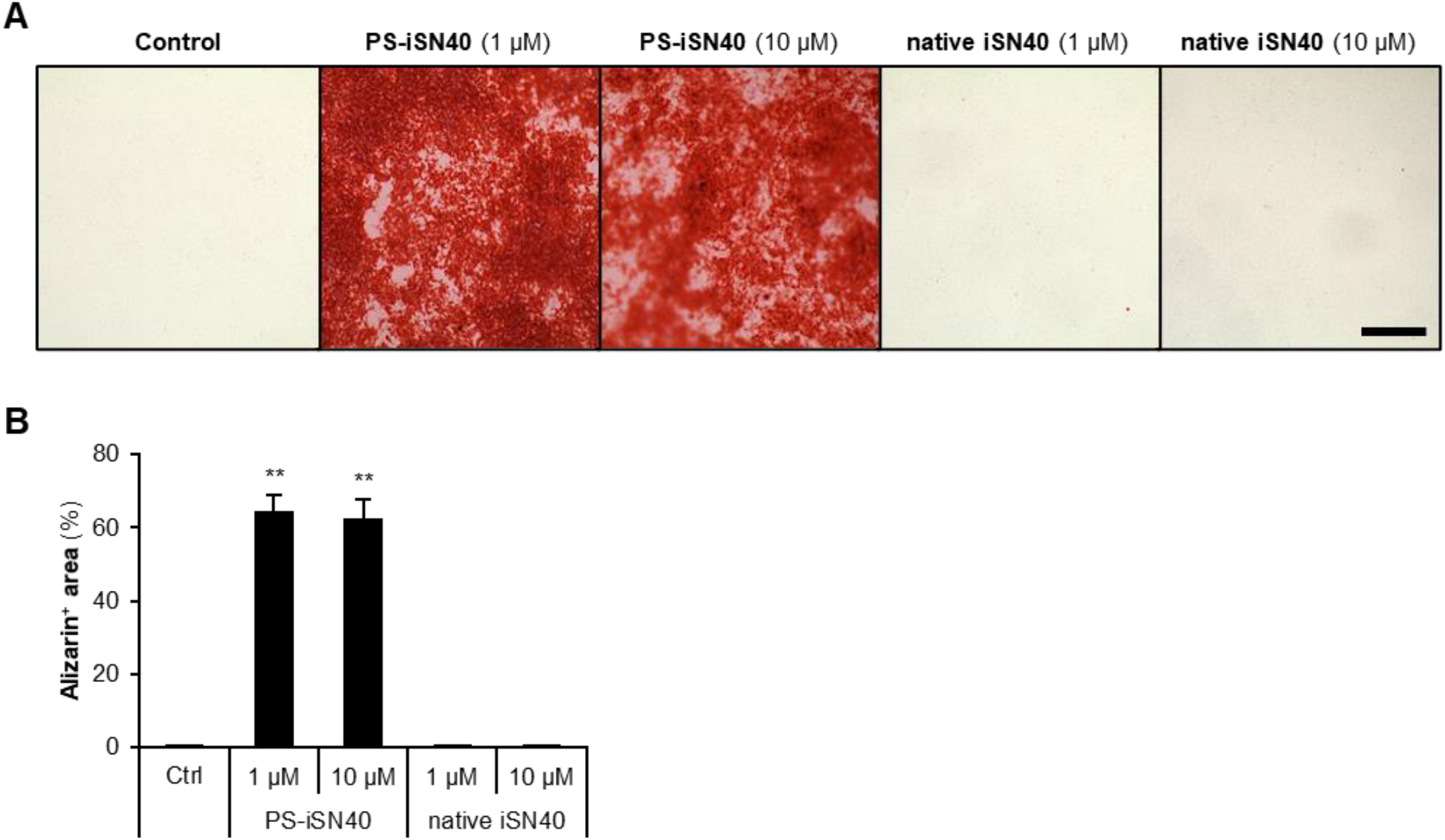
phosphorothioation is required for iSN40 to exert osteogenetic activity. (**A**) Representative images of alizarin staining of MC3T3-E1 cells treated with 1 or 10 μM of PS-iSN40 or native iSN40 in DM with 50 μg/ml AA for 10 days. Scale bar, 200 μm. (**B**) Quantification of alizarin-positive area. ** *p* < 0.01 vs control (Scheffe’s *F*-test). *n* = 4.

**Supplementary Figure S3.**
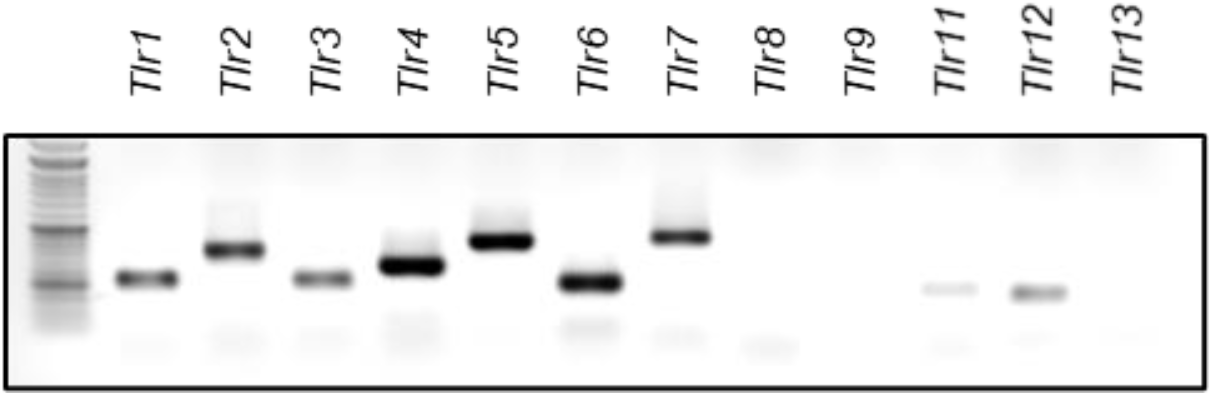
Expression of TLR genes in MC3T3-E1 cells. Total RNA of MC3T3-E1 cells maintained in GM was subjected to RT-PCR (40 cycles). Then the PCR products of TLR genes were subjected to agarose gel electrophoresis.

**Supplementary Table S1.**
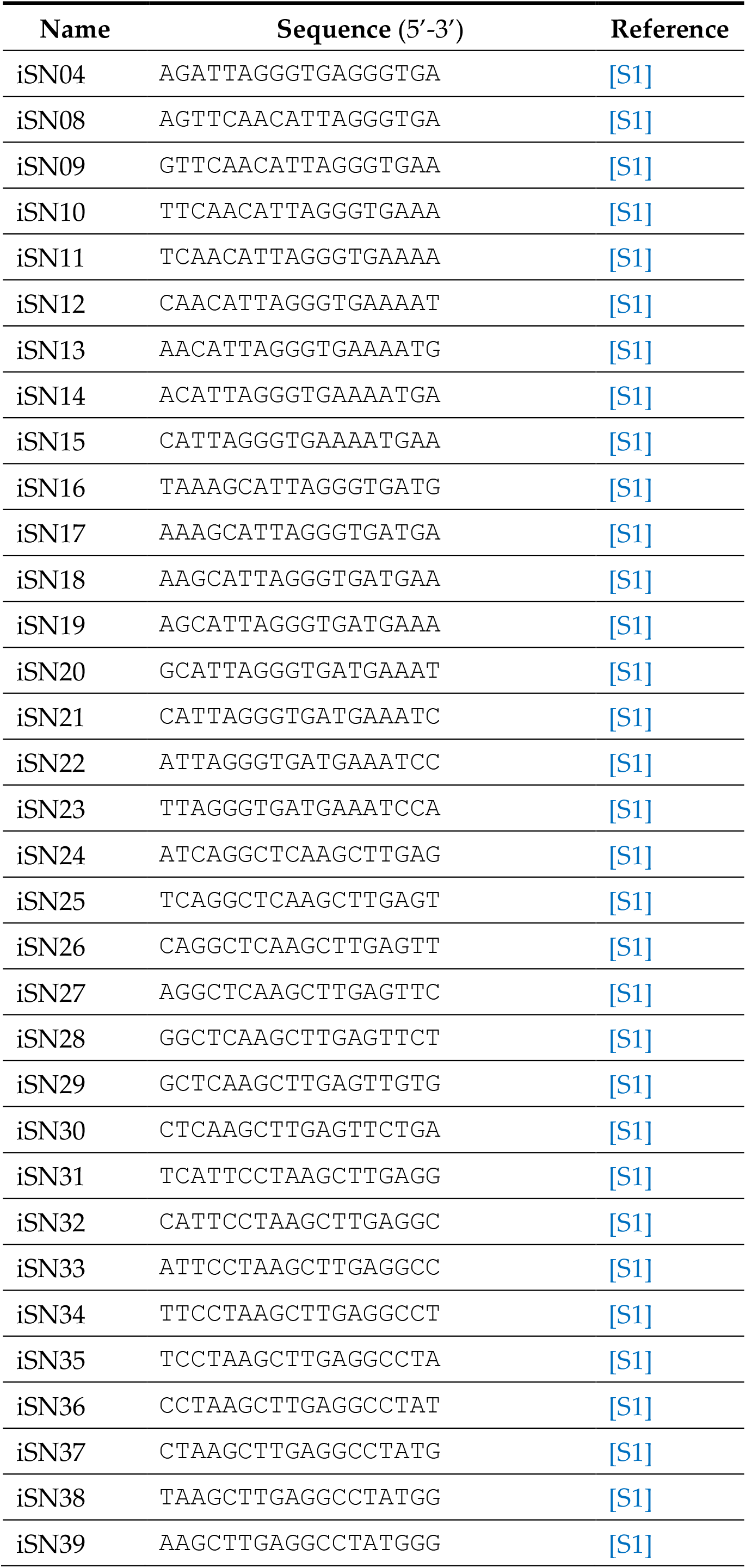
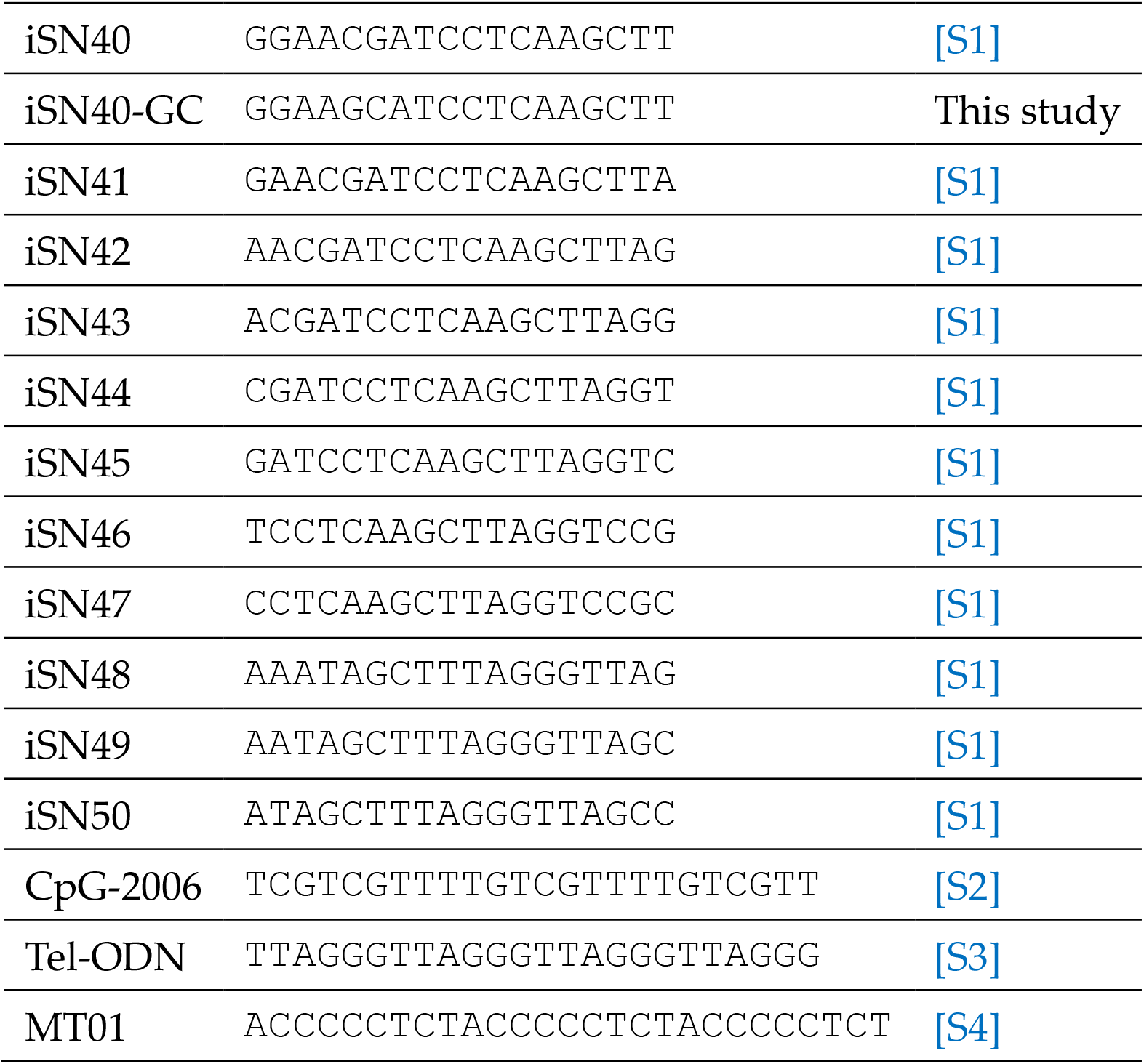
ODN sequences.

**Supplementary Table S2.**
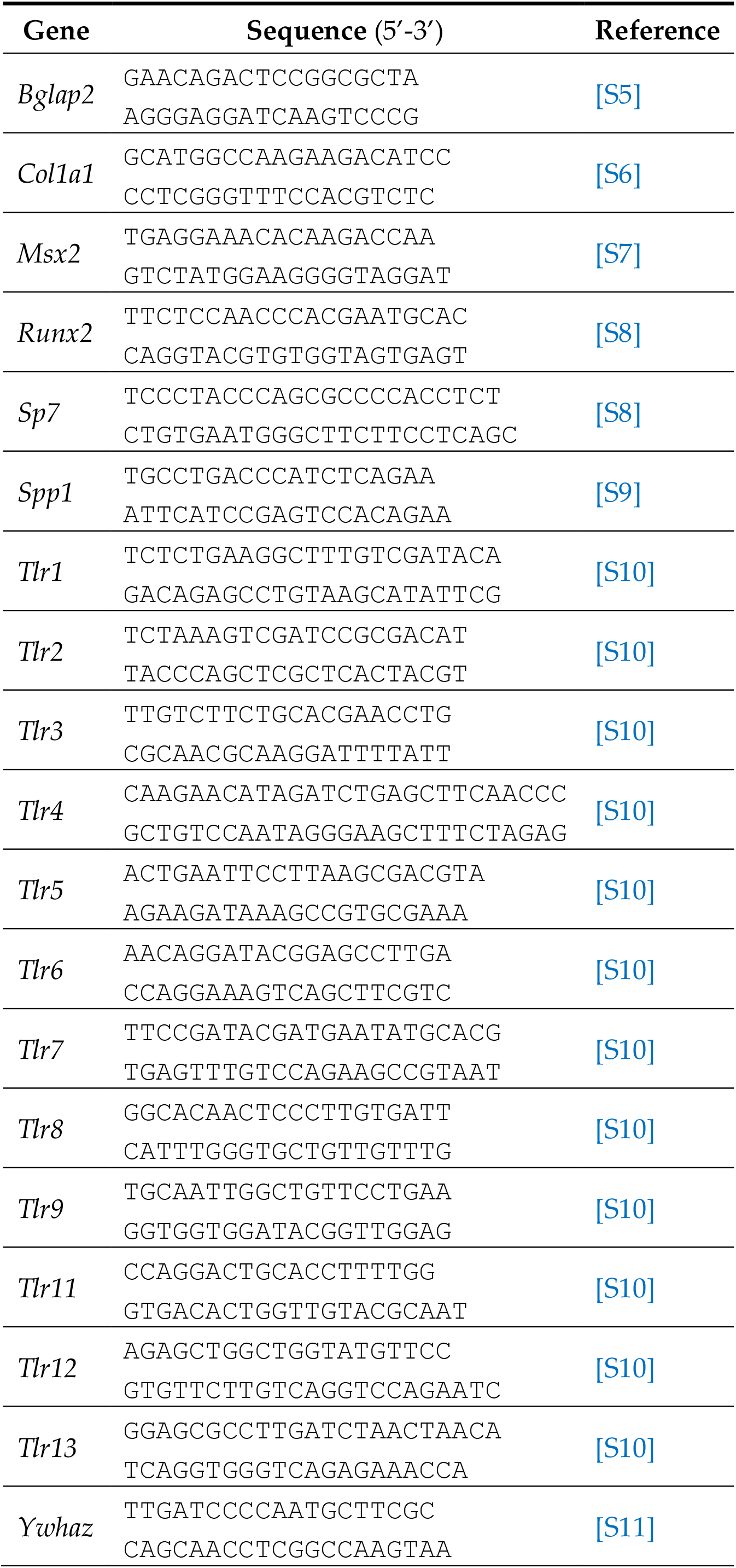
Primer sequences for RT-PCR.

